# Addressing the Most Neglected Diseases through an Open Research Model: the Discovery of Fenarimols as Novel Drug Candidates for Eumycetoma

**DOI:** 10.1101/258905

**Authors:** Wilson Lim, Youri Melse, Mickey Konings, Hung Phat Duong, Kimberly Eadie, Benoît Laleu, Ben Perry, Matthew H. Todd, Jean-Robert Ioset, Wendy W.J. van de Sande

## Abstract

Eumycetoma is a chronic infectious disease characterized by a large subcutaneous mass, often caused by the fungus *Madurella mycetomatis.* A combination of surgery and prolonged medication is needed to treat this infection with a success rate of only 30%. There is, therefore, an urgent need to find more effective drugs for the treatment of this disease. In this study, we screened 800 diverse drug-like molecules and identified 215 molecules that were active *in vitro.* Minimal inhibitory concentrations were determined for the 13 most active compounds. One of the most potent compounds, a fenarimol analogue for which a large analogue library is available, led to the screening of an additional 35 compounds for their *in vitro* activity against *M. mycetomatis* hyphae, rendering four further hit compounds. To assess the *in vivo* potency of these hit compounds, a *Galleria mellonella* larvae model infected with *M. mycetomatis* was used. Several of the compounds identified *in vitro* demonstrated promising efficacy *in vivo* in terms of prolonged larval survival and/or reduced fungal burden. The results presented in this paper are the starting point of an *Open Source Mycetoma (MycetOS)* approach in which members of the global scientific community are invited to participate and contribute as equal partners. We hope that this initiative, coupled with the promising new hits we have reported, will lead to progress in drug discovery for this most neglected of neglected tropical diseases.

**Author summary:** Mycetoma is a poverty-associated disease that was recently recognised as a neglected tropical disease by the World Health Organisation (WHO). This disease can be caused by either bacteria (actinomycetoma) or fungi (eumycetoma). The most common causative agent of mycetoma is the fungus *Madurella mycetomatis.* Actinomycetoma can be easily treated, but for eumycetoma, the current and only antifungal drug used is only able to successfully treat 30% of patients. Treatment often involves prolonged medication use and amputation of the affected area. This disease is disfiguring and is a social stigma for patients in endemic countries. To improve treatment for patients, we have looked at over 800 diverse drug-like molecules and compounds in hope to develop new drugs in this study. We have identified 215 compounds with activity against *M. mycetomatis* in vitro and several in vivo with our *Galleria mellonella* larvae model. We have chosen an open source approach with this study and placed our findings in an online database and made it available to the public. We invite the global scientific community to participate in our study and contribute as equal partners as long as an open source approach is held in hopes to fast track and boost drug discovery for Eumycetoma.

## Introduction

Mycetoma is a chronic infectious and inflammatory disease, characterised by a large subcutaneous mass and the excretion of grains, usually affecting the lower limbs [1]. This infection can be caused by either bacteria (actinomycetoma) or fungi (eumycetoma) [1] and the most common causative agent of mycetoma is the fungus *Madurella mycetomatis* [2]. Given its diverse etiology, it is not surprising that the course of treatment depends on the causative agent [3]. In general, actinomycetoma can be treated with medication only, with success rates of up to 90% [4], while for eumycetoma a combination of surgery and prolonged medication is needed [3]. Ketoconazole has been the mainstay of medical treatment for decades, given in a dose of 400-800 mg/day for a year with 50-70% success rates when combined with surgery [5, 6]. Recently, its use was restricted by the European Medicines Agency (EMA), the United States Food and Drug Administration (FDA) and the government of Sudan due to potentially fatal liver injury, drug interactions and adrenal gland problems [7, 8]. This restriction causes a dilemma, as few other options are available. Itraconazole is the most widely used alternative and in a recent study no serious hepatoxicity was noted (400 mg/day for 3 months, then 200 mg/day for 9 months) [9]. In all patients, the lesions were reduced enabling less mutilating surgery but, as is the case for ketoconazole, the fungus was still viable when isolated from surgical material [9, 10]. Other azoles have been used to some extent and with some success, but these are usually too expensive for use in endemic regions. Indeed, the only affordable drug for endemic areas was ketoconazole, at a cost of $30/month. With an average monthly income of only $60/month, itraconazole at $330/month is already considered to be too expensive for the patient and the affordability problem becomes more acute with the newer generation of antifungal agents such as voriconazole and posaconazole [11].

There is an urgent need to find an effective, safe and affordable oral antifungal agent with a short treatment duration for eumycetoma. Traditionally, the pharmaceutical industry has taken the lead in the development of novel antimicrobial drugs, but antimicrobial drug development programs have been drastically reduced in recent decades due to increased costs, decreased return on investment and the prospect of a long and expensive development process. Furthermore, drugs have never been specifically developed for neglected infectious tropical diseases such as mycetoma, due to the lack of sufficient potential return on investment. Therefore, in order to find a new, safe and effective treatment for eumycetoma, alternative approaches are necessary. A strategy that has gained much attention recently is to screen drugs already approved for other indications or drug candidates with a historical track record of development. This approach, also known as drug repurposing, is appealing as it is expected to reduce the overall costs and the timeframe required for drug development as well as de-risking the development process by capitalizing on preclinical and possibly phase I clinical data packages already available from a pre-existing drug development program [12].

The Pathogen Box is a set of 400 diverse, cherry-picked drug-like molecules previously shown to be active against pathogens causing tropical and neglected diseases. The Box includes a collection of approved drugs – the so-called reference set – used for the treatment of such diseases [13] and is made available free of charge by the Medicines for Malaria Venture (MMV) as an Open Access initiative tool to stimulate research and development on neglected diseases [12, 13]. In return, researchers are asked to share any data generated in the public domain within 2 years, creating an open and collaborative forum for infectious disease drug research. The chemical structures of the molecules in the Pathogen Box are publicly available and all compounds are annotated with valuable biological data sets arising from phenotypic screens, including cytotoxicity. Additionally, key *in vitro* and *in vivo* data related to drug metabolism and pharmacokinetics (DMPK) have recently been associated with all 400 compounds. A related MMV drug repurposing subset, the Stasis Box, has also been made available and consists of 400 compounds selected by medicinal chemistry experts out of 8000 compounds which have entered preclinical or clinical development but have been discontinued for various reasons, for example a lack of efficacy [14]. All compounds included in the Stasis Box can be purchased from commercial suppliers and have their chemical structures available in the public domain.

A screen of diverse agrochemicals against the pathogen causing Chagas disease, *Trypanosoma cruzi*, identified the fungicide fenarimol as an interesting starting point for potential therapeutics [15]. A collection of over 800 compounds was synthesized during a lead optimization program aimed at improving activity against *T. cruzi* [15–17] and addressing, through chemical modification including scaffold-hopping, the few identified liabilities of the fenarimol series notably low solubility, high *in vitro* metabolism and inhibition of cytochrome P450 3A4/5 (CYP3A4/5). This work resulted in the identification of a couple of optimized leads associated with a CYP51 selective profile (i.e. lacking cytochrome P450 3A4 inhibition) with nanomolar *in vitro* potency vs. *T. cruzi,* excellent *in vivo* efficacy in an infected murine model [15] and a lack of adverse events on extended dosing in murine exploratory toxicity studies. The extremely promising efficacy and drug-like characteristics of these two CYP51 inhibitors led to their further profiling with a view to potential development as new treatments for Chagas disease, but these efforts have been discontinued due to the clinical failure of azoles that act *via* an identical mechanism of action [18, 19].

Here we describe the screening of the 800 drug-like compounds contained in the Pathogen and Stasis Boxes, as well as selected compounds from the fenarimol-based chemical library, for their ability to inhibit the *in vitro* growth of *M. mycetomatis.* The hits with greatest potential were verified through chemical resynthesis and further evaluated in our grain model in *G. mellonella* larvae to determine the *in vivo* efficacy against *M. mycetomatis* grains. Finally, we propose a strategy for executing hit-to-lead campaigns based on these results that takes a highly collaborative, community-centered approach.

## Material and Methods

### Chemical libraries

The Pathogen and Stasis Boxes were kindly provided by Medicines for Malaria Venture (MMV, Geneva, Switzerland). The compounds were obtained in a 10 mM solution in dimethyl sulfoxide (DMSO) (Table S1). The fenarimol analogues were kindly provided by DND*i via* Epichem (Perth, Australia) and dissolved in DMSO, to reach a concentration of 10 mM. The resynthesis of hit compounds, and the chemical synthesis of the novel analogues reported, are described in S2 Text.

### Mycetoma strains

In this study, *M. mycetomatis* genome strain MM55 [20] was used to screen the Pathogen and Stasis Boxes and to determine the IC50 and IC90 of each of the compounds with inhibitory activity at a concentration of 100 µM. *In vivo* efficacy was also determined for this strain. To assess the activity of the selected compounds against other *M. mycetomatis* isolates, isolates P1, MM25, MM36, MM50, MM55, MM68 and MM83 were selected [21]. To assess the activity of the fenarimol analogues, strains MM13, MM14, MM26, MM30, MM41, MM45, MM49, MM50, MM54 and MM55 were selected. Strains MM26 and MM45 belonged to Amplified Fragment Length Polymorphism (AFLP) cluster I, strains P1, MM13, MM14, MM25, MM30, MM36, MM41, MM49, MM50, MM54, MM55 and MM68 belonged to AFLP cluster II and strain MM83 belonged to AFLP cluster III [21]. All MM-isolates have been isolated from patients diagnosed with mycetoma seen in the Mycetoma Research Centre, Khartoum, Sudan; the P1 strain originated from a patient in Mali [22]. The strains were isolated by direct culture of the black grains obtained by a deep surgical biopsy and were identified at the species level by morphology, polymerase chain reaction with *M. mycetomatis* specific primers [23] and sequencing of the internal transcribed spacers [24, 25]. The isolates were maintained in the laboratory on Sabouraud Dextrose Agar (Difco laboratories, Becton and Dickinson, Sparks, USA).

### Screening the Pathogen and Stasis Boxes

In order to screen the Pathogen and Stasis Boxes, an *M. mycetomatis* hyphal suspension was made as described previously [26]. In short, *M. mycetomatis* was cultured for 7 days at 37°C in RPMI 1640 medium supplemented with 0.3 g/L L-glutamine and 20 mM morpholinepropanesulfonic acid (MOPS). The mycelia were harvested by centrifugation and homogenized by sonication for 20 s at 28 µm (Soniprep, Beun de Ronde, The Netherlands). The fungal suspension was diluted in RPMI 1640 medium to obtain a transmission of 70% at 660 nm (Novaspec II, Pharmacia Biotech) [27]. In each well of a 96-well microplate, 100 µL of suspension was added. 1 µL of compound was added per well to obtain a final concentration of 100 µM. The microplate was taped to prevent evaporation and incubated for 7 days at 37°C. To facilitate end-point reading, 2,3-bis(2-methoxy-4-nitro-5-sulfophenyl)-5-[(phenylamino)carbonyl]-2H-tetrazolium hydroxide (XTT) was added to a final concentration of 25 µg/well and incubated for 2 h at 37°C, and another 3 h at room temperature [27, 28]. Plates were centrifuged and the extinction of 100 µL of supernatant was measured at 450 nm using the spectrophotometer. Complete growth reduction was defined as 80% or more reduction in viable fungal mass [26]. Each compound was tested in triplicate. To establish which compounds were most the potent in inhibiting *M. mycetomatis* growth, the concentration at which a 50% reduction in growth was obtained (IC50) was determined. In order to do this, all positive compounds were tested at concentrations of 0.098 µM, 0.39 µM, 1.56 µM, 6.25 µM and 25 µM. Growth was plotted against concentration and the IC50 was determined from the resulting graphs by visual reading. IC50s were determined in duplicate and the means plus standard deviations were determined in Excel. Four positive growth controls were included in each plate, consisting of a well in which the fungus was exposed to solvent only. Furthermore, two antifungal agents, namely posaconazole and amphotericin B, were included in the plates as positive controls for growth inhibition.

### Minimal inhibitory concentrations of seven *M. mycetomatis* strains

To determine if the most potent compounds identified above were also active against *M. mycetomatis* isolates with a different genetic background, seven *M. mycetomatis* isolates with different AFLP types were selected. Minimal inhibitory concentrations (MIC) were determined using the same protocol as described above. Concentrations ranged from 0.007 µM to 32 µM.

### Toxicity in *G. mellonella* larvae

To assess the toxicity of the compounds identified in the *in vitro* screenings *in vivo,* each compound was tested for toxicity in *G. mellonella* larvae. For each compound, a single dose was injected into the last right proleg with an insulin needle. Final concentrations of compound in each larva were 0.2 µM, 2 µM or 20 µM of compound. Controls were injected with distilled water only. Larval survival was monitored for 10 days. A non-significant difference in larval survival between the treated group and the control group indicated a lack of toxicity up to the dose under investigation.

### Infection of *G. mellonella* larvae with *M. mycetomatis* and antifungal treatment

*G. mellonella* larvae were infected with *M. mycetomatis* isolate Mm55 according to a previously published protocol [29]. In short, *M. mycetomatis* mycelia were cultured in colourless RPMI 1640 medium supplemented with L-glutamine (0.3 g/L), 20 mM morpholinepropanesulfonic acid (MOPS) and chloramphenicol (100 mg/L; Oxoid, Basingstoke, United Kingdom) for 2 weeks at 37°C and then sonicated for 2 min at 28 µm. The resulting homogenous suspension was washed once in PBS and further diluted to an inoculum size of 4 mg wet weight per larva. Inoculation was performed by injecting 40 µL of the fungal suspension into the last left pro-leg with an insulin 29G U-100 needle (BD diagnostics, Sparsk, USA). Larvae injected with PBS were included as controls. Larvae were treated with 20 µM of compound per larva or with solvent only. Each group consisted of 15 larvae, and each experiment was performed three times. The results of the three individual experiments were pooled and plotted in survival curves using GraphPad Prism 5. To treat the larvae, compounds were administered at 4, 28 and 52 hours after infection. Treatment was started at 4 hours since grains were already visible at that time point. To monitor the course of infection, larvae were checked daily for survival for 10 days. If during these 10 days larvae formed pupa, these individuals were not considered further, since we could not ascertain whether these individual larvae would have survived or died during the course of the infection. This means that within each treatment group, the maximum number of larvae was 45 (if no pupae were formed).

### Fungal burden

To determine the fungal burden, 5 larvae from each group were sacrificed at day 3 post infection. First, haemolymph was collected and measured as described earlier [29] to assess melanization. To assess the number of grains per larva, larvae were fixed in 10% buffered formalin and dissected longitudinally into two halves with a scalpel and processed for histology [30]. Sections were stained with haematoxylin and eosin (HE) and Grocott methanamine silver, and grains were manually counted under a light microscope mounted with a Canon EOS70D camera (Canon Inc) by two independent scientists. As controls, infected and non-infected larvae treated with PBS were used. Grains were magnified 40x and visualized on the computer screen using the supplied EOS Utility™ software (Canon Inc). Grains were categorized into large, medium or small sizes using the enlargement display frame present in the Live View Shooting mode. Under 40x magnification, the enlargement display frame has a width and height of approximately 250 µm and 160 µm and sums up to a dimension of 0.04 mm^2^. Grains that were larger than half of the display frame were categorized as large (>0.02 mm^2^). Grains that were larger than a quarter of the frame but smaller than half of the frame are categorized as medium (0.01 - 0.019 mm^2^) and those between one-eighth to a quarter of the display frame (0.005 - 0.009 mm^2^) were categorized as small. The sum of all large, medium and small grains present in larvae was used to represent the total number of grains in the larvae. To determine the total size of grains in the larvae, the sum of all grains in a larva multiplied by the minimum size of their respective category (large: 0.02mm^2^, medium: 0.01mm^2^ and small: 0.005mm^2^) was used.

### Statistical analysis

To compare survival curves, the Log-rank test was performed with GraphPad Prism 5 (version 5.03, GraphPad Inc.) To determine if there was a statistical difference across the different groups in terms of melanization and grain counts, the Kruskal-Wallis test was performed with GraphPad Prism 5. If a difference was found with the Kruskal-Wallis test, pair-wise comparisons were made between the PBS treated groups and the different antifungal treated groups with the Mann-Whitney U test with GraphPad Prism 5 to determine differences in melanization or CFU count. A p-value smaller than 0.05 was deemed significant. All negative values after normalization were refigured to zero for statistical analysis. To determine the statistical difference in the total number and sizes of grains between the treated and non-treated groups, a Mann-Whitney test was performed with GraphPad Prism 5. A p-value smaller than 0.05 was deemed significant.

## Results

### *In vitro* screening of the Pathogen and Stasis Boxes identify MMV006357 and MMV689244 as the most potent exploratory compounds

In total, 800 different drug-like compounds originating from either the Pathogen or Stasis Boxes were evaluated for their potential to inhibit the growth of *M. mycetomatis in vitro* and *in vivo* using a sequential assay workflow (Figure 1). Of the 800 compounds screened at a concentration of 100 µM, 215 compounds inhibited growth of *M. mycetomatis.* Of these 215 compounds, 147 originated from the Pathogen Box and 68 from the Stasis Box (Figures 1, 2A and S1 Table). A significantly higher hit rate was obtained with the Pathogen Box (36.8%) than with the Stasis Box (17.0%) (Fisher Exact, p<0.0001). To determine which of these compounds were most potent at inhibiting the growth of *M. mycetomatis,* the IC50 values of these 215 compounds were determined (S1 Table, Figure 2B). It appeared that the median IC50 of these 250 compounds was 47.8 µM (<0.09 – 77.5). In total, 13 compounds had an IC50 of 5 µM or lower (Table 1). Twelve compounds originated from the Pathogen Box and one from the Stasis Box. Surprisingly, the antifungal agent amphotericin B, also present as a reference compound in the Pathogen Box, had an IC50 of only 9.7 µM and was therefore excluded from further evaluation; this compound had previously been demonstrated to be active against *M. mycetomatis* both *in vitro* [26] and *in vivo* [31]. To determine if the 13 most potent compounds were also able to inhibit *M. mycetomatis* isolates with a different genetic background or origin, six other *M. mycetomatis* isolates were selected based on their AFLP type or origin and MICs were determined for all 13 compounds. As can be seen in Table 1, Figure 3 and S1 Table, MMV688774 (posaconazole, a triazole-based drug approved for the treatment of fungal infections in humans) was the most potent inhibitor of *M. mycetomatis* growth with an MIC50 of <0.007 µM. Two azoles, MMV688942 (bitertanol) and MMV688943 (difenoconazole), and two strobilurins, MMV021057 (azoxystrobin) and MMV688754 (trifloxystrobin) – all 4 products being used as agrochemicals – also had strong activity against *M. mycetomatis* with MIC50s ranging from 0.06 µM to 0.125 µM. The remaining 8 hits – all part of the non-reference set of exploratory compounds – displayed lower potencies against *M. mycetomatis* with MIC50 values ranging from 0.25 µM to 8 µM. Compounds MMV006357 and MMV689244 were the most potent of these 8, displaying MIC50 values of 0.25 µM and 1 µM, respectively. Of note, compound MMV689244 is a fenarimol analogue, identified as a potent inhibitor of *Trypsonosoma cruzi* during a targeted screening exercise for new drugs for Chagas disease [15–17], during which over 800 fenarimol analogues were synthesized (MMV689244 corresponds to EPL-BS1246 in this library).

**Fig 1.**
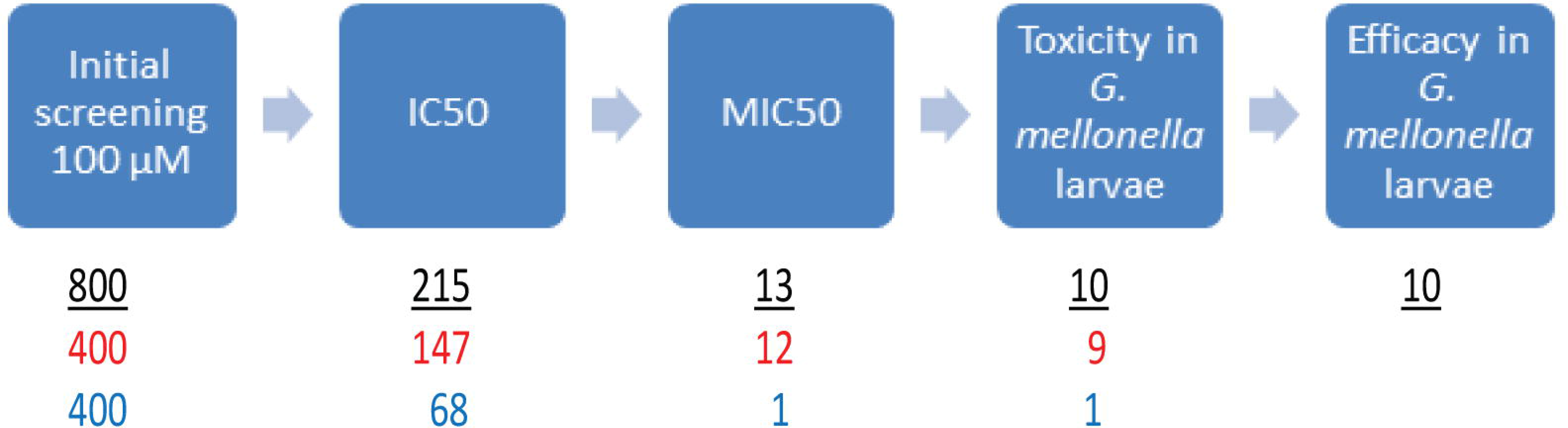
Flow diagram for *in vitro* and *in vivo* evaluation of all compounds (red and blue numbers show numbers of hits from the Pathogen and Stasis Boxes respectively, with the black numbers indicating totals).

**Fig 2.**
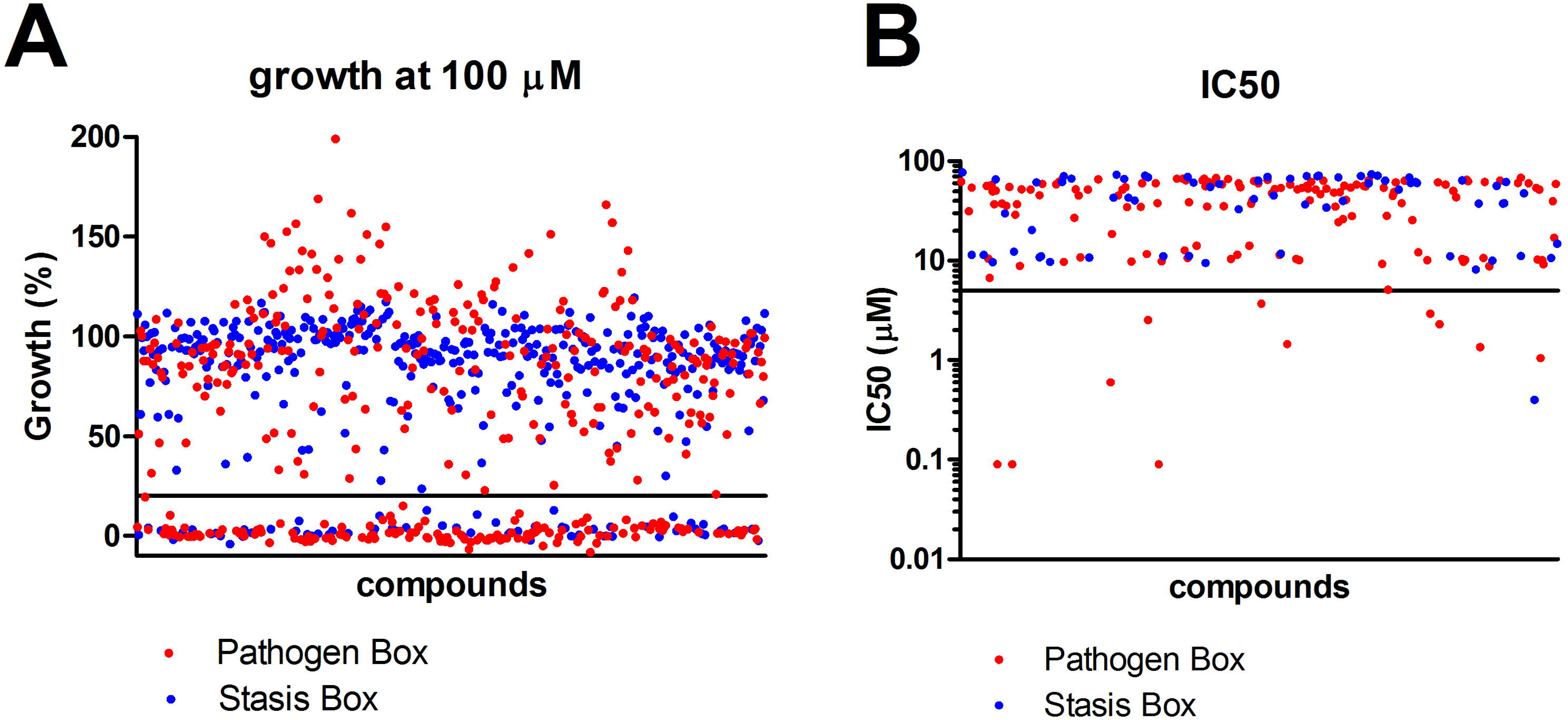
Percentage growth at 100 µM (panel A) and IC50 (panel B) of selected compounds.

**Fig 3.**
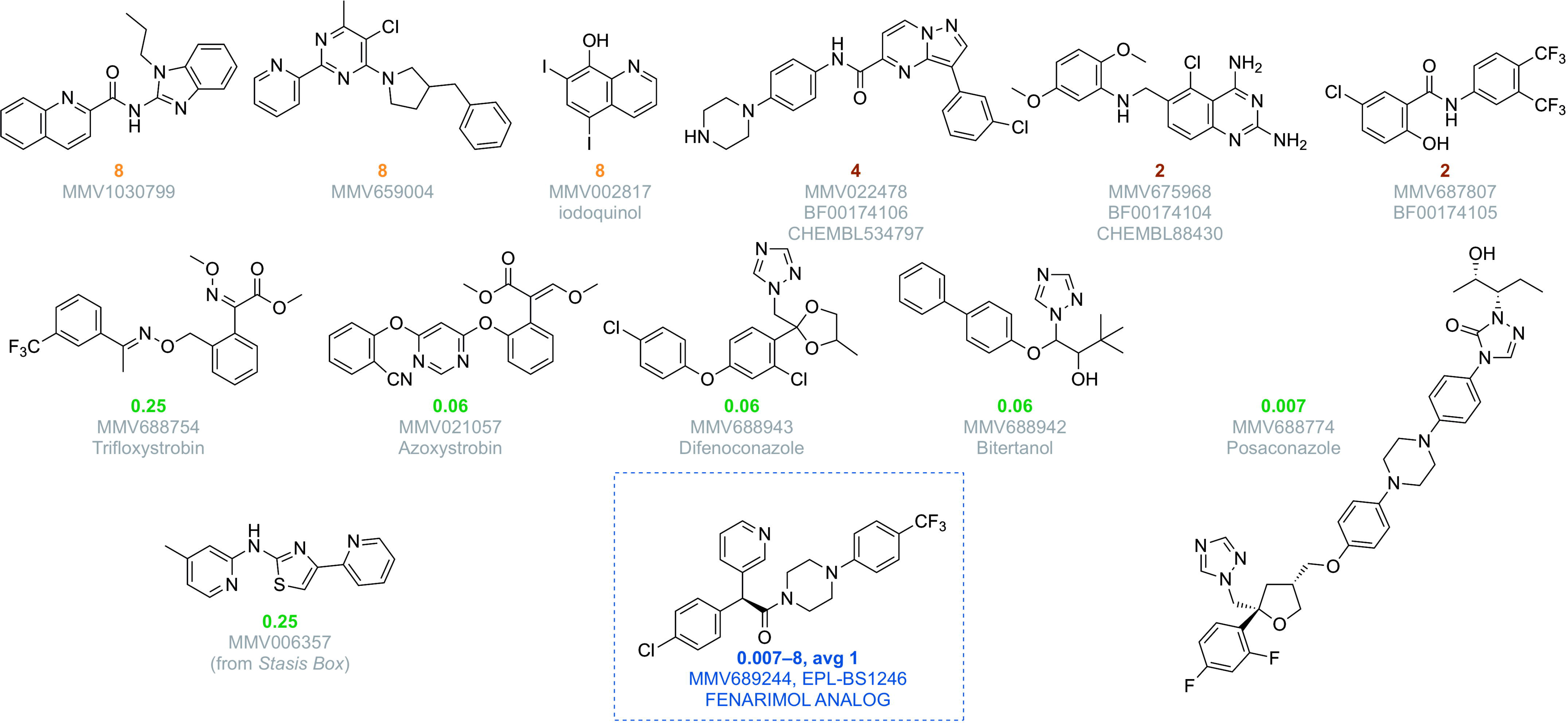
MIC50 in micromolar obtained after testing each of the compounds against seven different *M. mycetomatis* isolates.

**Table 1.** IC50 and MIC50 against *M. mycetomatis* of the 13 most active compounds identified in the Pathogen and Stasis Boxes

|  | Trivial name or<br>ChEMBL code | Class | MoA | use | IC50 of<br>strain<br>MM55 | MIC50 |
| --- | --- | --- | --- | --- | --- | --- |
| **MMV688774** | Posaconazole | Azoles | CYP51 inhibitor | antifungal (human) | <0.10 | <0.007 |
| **MMV688942** | Bitertanol | Azoles | CYP51 inhibitor | antifungal (agrochemical) | <0.10 | 0.06 |
| **MMV688943** | Difenoconazol | Azoles | CYP51 inhibitor | antifungal (agrochemical) | <0.10 | 0.06 |
| **MMV021057** | Azoxystrobin | Strobilurins | Complex III mitochondrial<br>electron transport chain binder | antifungal (agrochemical) | 0.60 | 0.06 |
| **MMV688754** | Trifloxystrobin | Strobilurins | Complex III mitochondrial<br>electron transport chain binder | antifungal (agrochemical) | 1.05 | 0.25 |
| **MMV006357** | N/A | 2-aminothiazole | N/A | N/A | 0.40 | 0.25 |
| **MMV689244** | EPL-BS1246 | Fenarimols (non-azoles) | CYP51 inhibitor | antiprotozoal (Chagas<br>disease preclinical<br>candidate) | 1.35 | 1 |
| **MMV687807** | N/A | Benzamides | N/A | N/A | 1.45 | 2 |
| **MMV675968** | ChEMBL88430 | Quinazolines | Dihydrofolate reductase | N/A | 2.30 | 2 |
| **MMV022478** | ChEMBL534797 | Pyrazolo-pyrimidines | N/A | N/A | 2.95 | 4 |
| **MMV002817** | Iodoquinol | Quinoline | Unknown | antiprotozoal (amoebiasis) | 2.55 | 8 |
| **MMV1030799** | N/A | Benzimidazole quinoline | N/A | N/A | 3.70 | 8 |
| **MMV659004** | N/A | Pyridyl-pyrimidine | N/A | N/A | 5.10 | 8 |

### *In vitro* screening of the 35 MMV689244 analogues identifies four potent fenarimol analogues

Due to the potent inhibition of *M. mycetomatis* by MMV689244, as well as the availability of the compound collection to our screening program, an additional 35 diverse compounds from the original 800 fenarimol analogue set were tested to determine their *in vitro* activity against *M. mycetomatis* hyphae (S1 Table and Figure 4). These compounds were selected based on similarity to the original hit whilst ensuring an even sampling of the various alternative scaffolds within the chemical set (Figure 5). The fenarimol analogues tested included variations of substituents around the fenarimol core as well as compounds with closely related scaffolds, since MMV689244 itself is a scaffold variant of fenarimol. Fenarimol analogues were tested for potential inhibition of the growth of *M. mycetomatis* at both 100 and 25 µM and the most potent compounds were then subjected to dose response determinations. Four of the analogues demonstrated inhibition of growth at both concentrations (Figure 4, top): EPL-BS0495, EPL-BS0800, EPL-BS1025 (racemate of EPL-BS1246) (all having MIC50 values of 4 µM) and EPL-BS0178 (MIC50 of 8 µM) (Figure 4). Interestingly, the potency of these analogues was not confined to a single scaffold type, with different representatives of the fenarimol core and fenarimol inspired scaffold changes resulting in MIC50 values in the micromolar (2-8 µM) range (Figure 4). In addition, EPL-BS1025 demonstrated similar potency to its enantiopure counterpart from the original assay (EPL-BS1246, MMV689244), validating this hit from the original screen.

**Fig 4.**
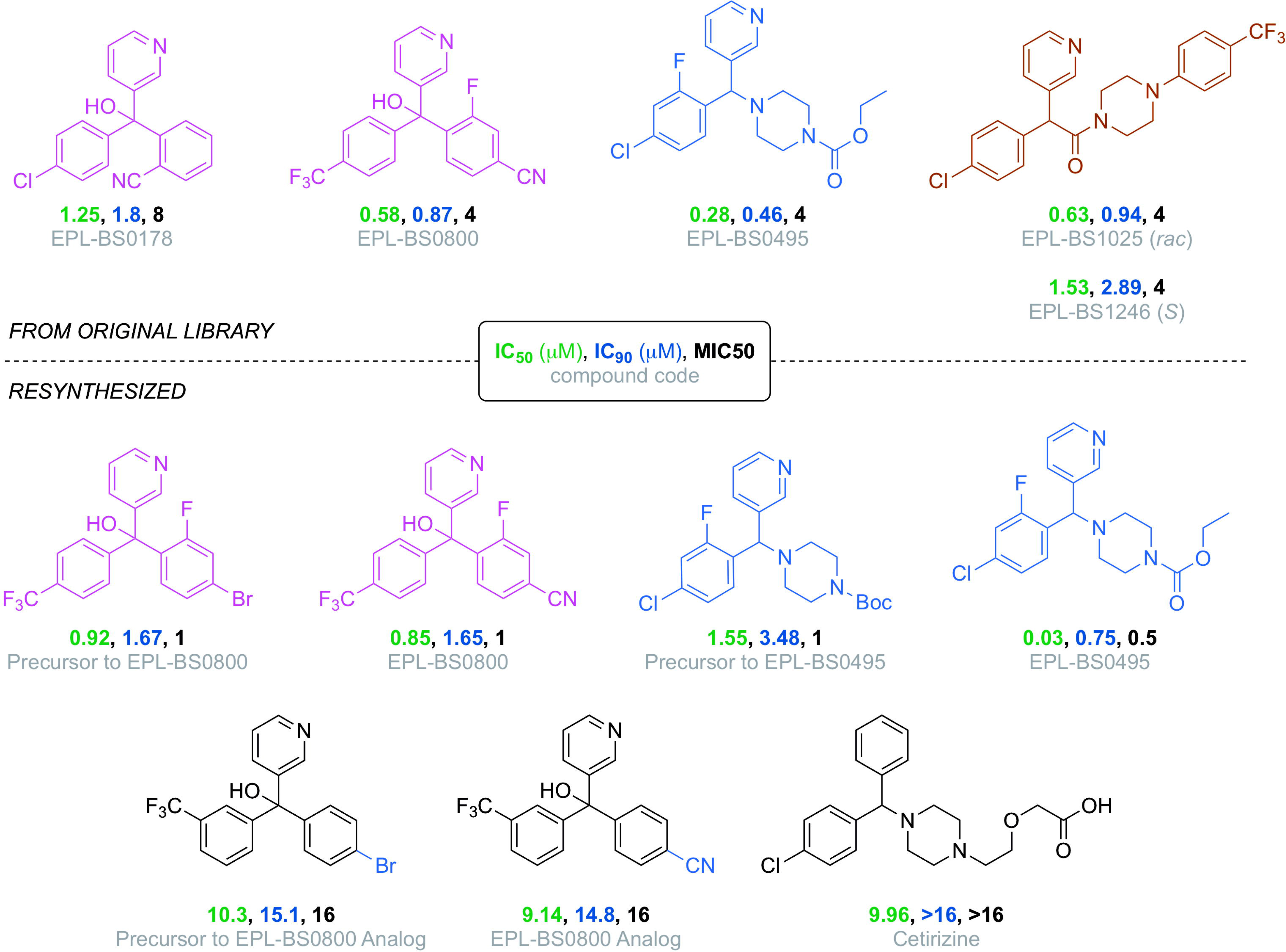
*In vitro* potency of the active compounds from the fenarimol library. Originally-evaluated samples (top); resynthesized samples and analogs (bottom).

**Fig 5.**
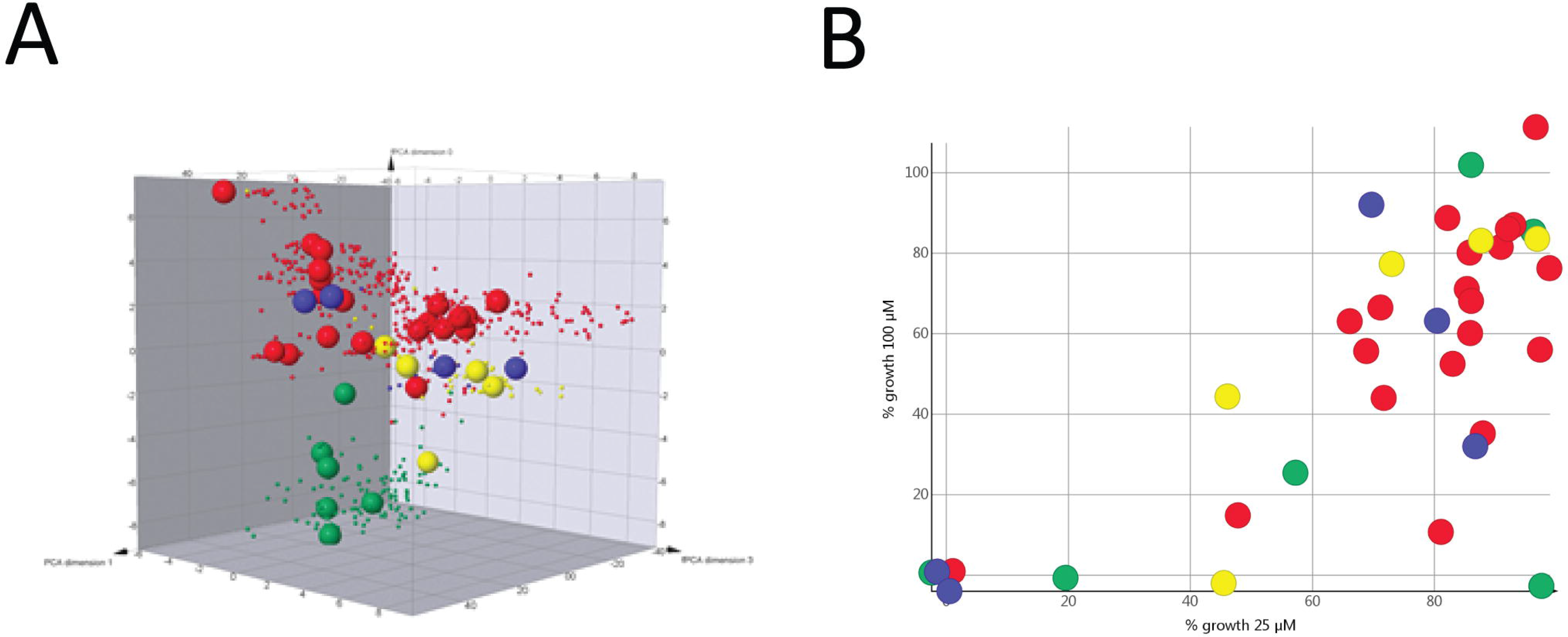
Chemical space representation of the 800 member fenarimol analogue library. A: Compounds chosen for testing are represented by oversized points, and different core scaffolds represented by colour (Blue-scaffold A; green scaffold B, red scaffold C, yellow scaffold D). B: % Growth inhibition by selected fenarimol analogues at 25 and 100 μM

Resynthesis of the fenarimol analogs was achieved using methods largely based on procedures found in the literature [15–17] as described in the supporting information (S2 Text), and these compounds were validated as hits (Figure 4, bottom). The resynthesized EPL-BS0800 and EPL-BS0495 showed potent *in vitro* activity, with a slightly higher IC50 for the resynthesized EPL-BS0800 (0.85 vs 0.58) and a slightly higher IC50 for the resynthesized EPL-BS0495 (0.03 vs 0.28) compared to the original compounds. Several synthetic precursors and a novel analog synthesized with minor changes in the pendant rings of EPL-BS0800 were also evaluated. While the novel analog of EPL-BS0800 did not provide *in vitro* potency, the precursors to EPL-BS0800 itself (containing a bromine atom in place of a cyano group) and to EPL-BS0495 (containing a Boc-protecting group rather than the ethyl carbamate) were active. These results suggest that either the carbamate group in EPL-BS0495 is labile under the assay conditions or there is room for tolerance to variation in this position on the molecule. The commercially available anti-inflammatory compound cetirizine (Zyrtec™), which possesses striking structural similarity to the fenarimol analogs, was evaluated but found to be inactive.

### *In vivo* tolerability of the 10 most potent compounds identified from the Pathogen and Stasis Boxes

The 10 most potent hits resulting from screening the Pathogen and Stasis Boxes were further evaluated *in vivo* using the *G. mellonella* larvae model. A first requirement was to determine if these compounds displayed any toxicity to the larvae. This was assessed by injecting a single dose of compound into the hemolymph of the larvae. Survival was monitored for 10 days. At the highest concentration tested (20 µM/larvae), none of the compounds displayed toxicity.

### *In vivo* activity of the 10 most potent compounds identified from the Pathogen and Stasis Boxes

Since none of the compounds were considered toxic, therapeutic efficacy was determined in *M. mycetomatis* infected larvae. Of the reference compounds used, only MMV688774 (posaconazole) (Log-Rank, p=0.011) and MMV688942 (bitertanol) (Log-Rank, p=0.0178) prolonged survival in this model in a statistically significant manner (Figure 6A). MMV688943 (difenconazole), MMV021057 (azoxystrobin) and MMV688754 (trifloxystrobin) did not prolong larval survival (Figures 6A and 6B). In *M. mycetomatis* infected larvae, grains are visible within 4 hours after infection and after three days they are fully matured. Within each of the larvae, grains of different sizes were noted, and these grains were classified as large, medium and small. After treating the larvae for three days with PBS or any of the compounds, significant differences in the total number of grains (large, medium and small together) were noted between PBS treated larvae and larvae treated with any of the tested compounds (Table 2, Figure 7A, Figure 8). A significant reduction in the number of grains was noted for MMV688942 (bitertanol) (Mann-Whitney, p=0.007), MMV688943 (difenconazole) (Mann-Whitney, p=0.002), MMV688774 (posaconazole) (Mann-Whitney, p=0.004), MMV021057 (azoxystrobin) (Mann-Whitney, p=0.029) and MMV688754 (trifloxystrobin) (Mann-Whitney, p=0.001) (Figure 7A). No difference in the distribution of large, medium and small grains was noted for any of the groups, except for MMV688942 (bitertanol) treated larvae (Table 2, Figure 8C). In that group, a significantly lower number of large grains and a higher number of smaller grains was noted compared to the PBS treated larvae. As a result, in addition to the total grain number, the total grain mass also differed significantly between PBS and compound treated larvae. The total grain mass obtained in PBS treated larvae was 0.08 mm^2^, which was significantly higher than the mass obtained for the majority of the compounds evaluated (Figure 7C). Since melanization of the haemolymph is part of the immune system of *G. mellonella* larvae, a difference in melanization is an indication of the activity of the larval immune system. A significant decrease in melanization was observed in MMV688943 (difenconazole) (Mann-Whitney, p=0.047) treated larvae, whereas no significant difference in melanization was observed with other compounds (Figure 7E). Of the five other compounds tested, only MMV006357 (Log-Rank, p=0.0012), MMV675968 (p<0.0001) and MMV022478 (p=0.0224) prolonged survival in a statistically significant manner (Figures 6C and 6D). The highest overall survival was obtained with compound MMV006357, which resulted in an overall survival of 28.6%. Compounds MMV675968 and MMV022478 demonstrated a lower overall survival percentage, but prolonged survival more effectively at the beginning of the infection. Of these compounds, only the fenarimol analogue MMV689244 (EPL-BS1246) was able to reduce the number of grains and the total grain mass produced significantly (Table 2, Figure 7A, C, Figure 8D). No difference in the distribution of large, medium and small grains was noted (Table 2). No significant difference in grain number, grain mass, grain distribution or melanization was observed for MMV006357, MMV675968, MMV687807 and MMV022478 (Table 2, Figure 7A, C, E).

**Fig 6.**
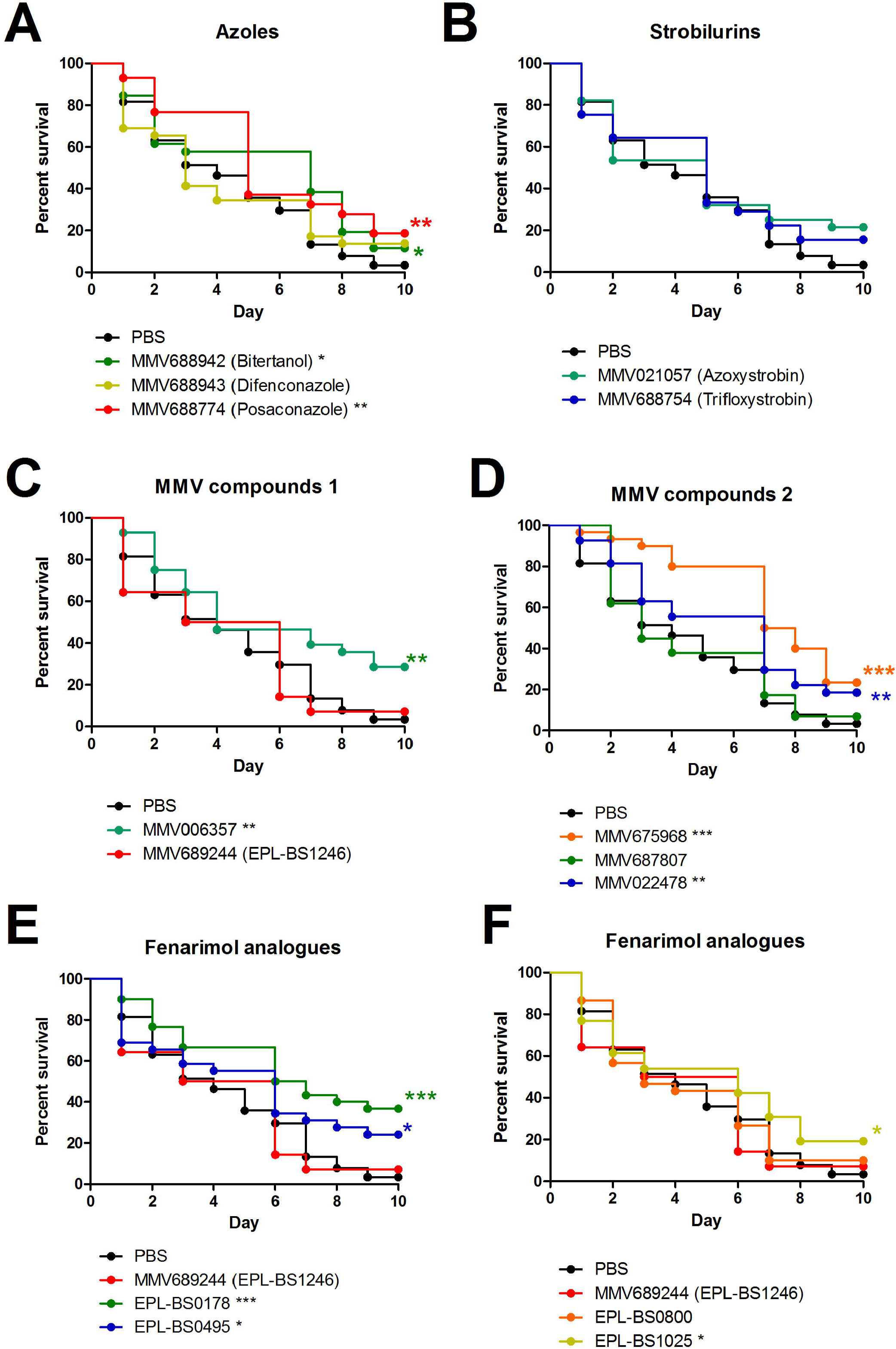
Survival curves of larvae infected with *M. mycetomatis* and treated with selected compounds. The black line in all panels corresponds to *M. mycetomatis* infected larvae treated with PBS. This is the control line. In panel A, the survival of larvae treated with azoles MMV688942 (Bitertanol), MMV68893 (Difenconazole) and MMV688774 (Posaconazole) is displayed. In panel B, the survival of larvae treated with strobilurins MMV021057 (azoxystrobin) and MMV688754 (trifloxystrobin) is displayed. In panels C and D, the survival of larvae treated with the MMV compounds is displayed. These include MMV006357, MMV6894244 (EPL-BS1246), MMV675968, MMV687807 and MMV022478. In panels E and F, the fenarimol analogues MMV689244 (EPL-BS1246), EPL-BS0178, EPL-BS0495, EPL-BS0800 and EPL-BS1025 are displayed. Significant survival was displayed as * (0.01<p<0.05), ** (0.00<p<0.01) or *** (0.001<p).

**Fig 7.**
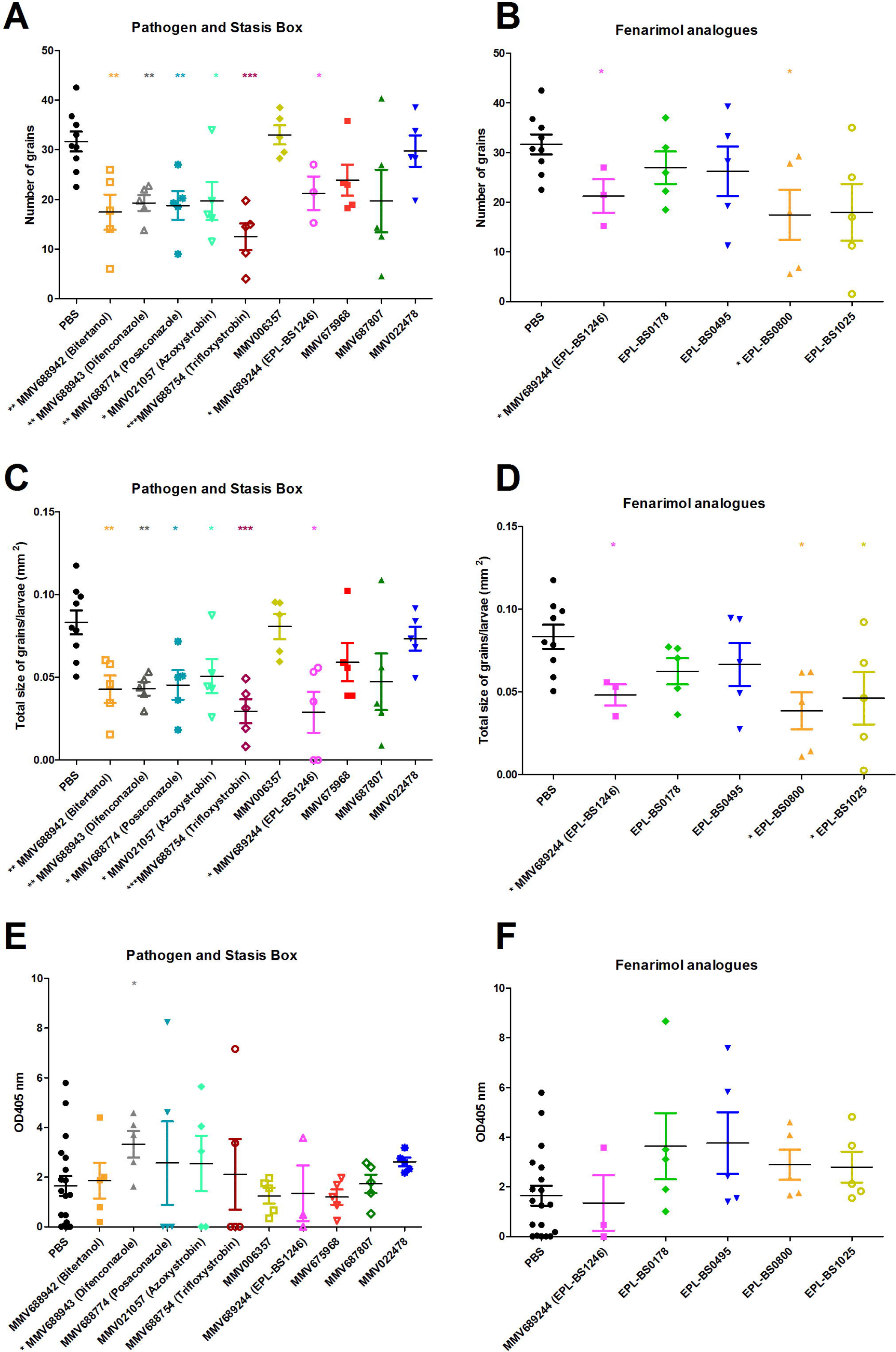
Number of grains (panels A and B), total grain size (panels C and D) and melanization (panels E and F) of larvae infected with *M. mycetomatis* and treated with selected compounds. PBS in all panels corresponds to *M. mycetomatis* infected larvae, treated with PBS. This is the control group. In panel A, C and E, compounds from the Pathogen and Stasis Box are displayed. In panel B, D and F, fenarimol analogues are displayed. Significant differences determined using the Mann-Whitney U-test were displayed as * (0.01<p<0.05), ** (0.001<p<0.01), or *** (p<0.001).

**Fig 8.**
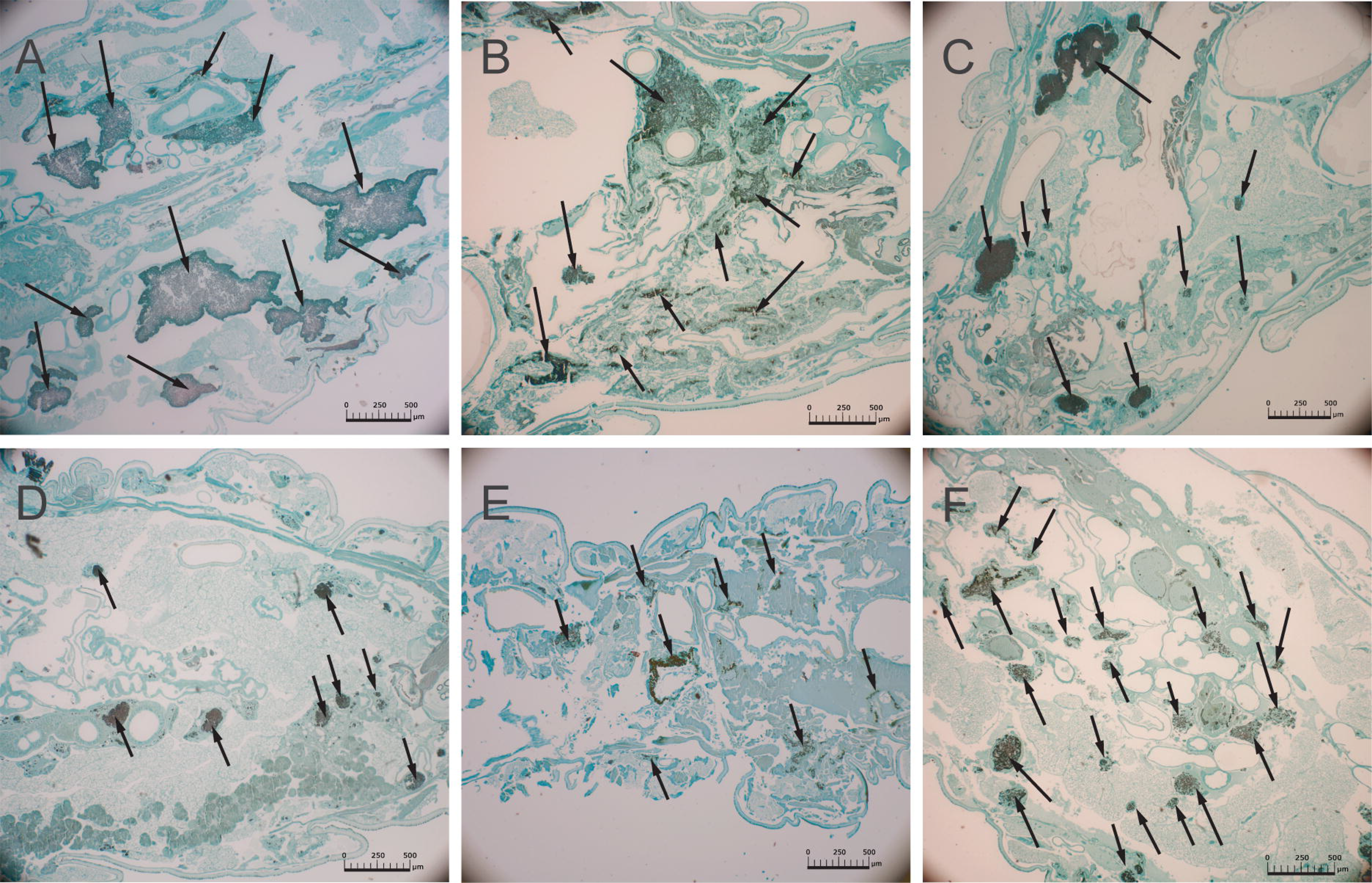
Histopathological sections of *Madurella mycetomatis* grains in infected *Galleria mellonela* larvae, treated with different compounds from the Pathogen and Stasis Boxes or with fenarimol analogues and sacrificed 72h after inoculation. Histopathological sections are stained with Grocott to demonstrate the presence of fungal grains (black stained) and indicated by arrows. Panels A and B demonstrate larvae treated with PBS as a control, for which both large grains (A) and smaller grains (D) are visible. Panels C and D show *G. mellonella* infected with *M. mycetomatis* and treated with MMV688942 (Bitertanol)(C), MMV689244 (EPL-BS1246) (D), EPL-BS0178 (E) and EPL-BS0495 (F). The scale bar present on each image represents 500µm.

**Table 2:** Statistical analysis of the total number of grains in larvae treated with the 14 compounds.

| Compounds | n | Median |  |  | Median of total grain number | Total number |  |  | p-value (Chi-square) |
| --- | --- | --- | --- | --- | --- | --- | --- | --- | --- |
|  |  | Large<br>n(STDEV) | Medium<br>n(STDEV) | Small<br>n(STDEV) |  | Large | Medium | Small |  |
| **Control** |  |  |  |  |  |  |  |  |  |
| PBS | 9 | 2.8 (2.1) | 8.5 (3.2) | 21.8 (3.6) | 31 | 32 | 68 | 185 | - |
| **Pathogen and Stasis Box** |  |  |  |  |  |  |  |  |  |
| MMV688942 (Bitertanol) | 5 | 1.5 (0.8) | 3 (1.6) | 12.5 (6.0) | 18 | 8 | 17 | 62 | 0.547 |
| MMV688943 (Difenconazole) * | 5 | 1.5 (0.9) | 2.8 (0.9) | 15.5 (3.2) | 20 | 6 | 15 | 76 | 0.047 * |
| MMV688774 (Posaconazole) | 5 | 1.8 (1.5) | 3.3 (1.9) | 13.5 (4.5) | 19 | 9 | 17 | 69 | 0.373 |
| MMV021057 (Azoxystrobin) | 5 | 1.8 (1.3) | 4.5 (1.5) | 10.3 (6.1) | 17 | 10 | 23 | 65 | 0.953 |
| MMV688754 (Trifloxystrobin) | 5 | 0.3 (1.3) | 2.3 (1.2) | 10.3 (4.4) | 15 | 5 | 11 | 46 | 0.373 |
| MMV006357 | 5 | 3 (2) | 6 (1.8) | 22.5 (2.8) | 33 | 15 | 33 | 117 | 0.425 |
| MMV689244 (EPL-BS1246) | 3 | 1.3 (1) | 4.3 (1.5) | 14 (5.7) | 22 | 4 | 12 | 48 | 0.266 |
| MMV675968 | 5 | 2 (2) | 4.5 (2.4) | 15.8 (2.6) | 23 | 11 | 26 | 83 | 0.689 |
| MMV687807 | 5 | 1.2 (1.9) | 2.5 (3.7) | 10.5 (9.1) | 14 | 8 | 18 | 72 | 0.297 |
| MMV022478 | 5 | 3 (0.9) | 5.5 (1.2) | 21 (5.7) | 29 | 14 | 29 | 106 | 0.423 |
| **Fenarimol analogues** |  |  |  |  |  |  |  |  |  |
| EPL-BS0178 | 5 | 1.3 (1) | 7 (2.5) | 15.8 (5.9) | 26 | 7 | 32 | 96 | 0.127 |
| EPL-BS0495 | 5 | 2.5 (1.8) | 5 (2.1) | 20.8 (7.9) | 28 | 16 | 21 | 94 | 0.195 |
| EPL-BS0800 | 5 | 1.3 (0.9) | 2.5 (1.6) | 12 (9.4) | 18 | 5 | 14 | 68 | 0.062 |
| EPL-BS1025 | 5 | 2.8 (1.6) | 2.5 (4.5) | 11.8 (7.3) | 17 | 10 | 20 | 60 | 0.946 |

### *In vivo* activity of the fenarimol analogues

One of the five non-reference compounds, MMV689244 (EPL-BS1246), did not prolong larval survival (Figure 6C), although it was able to reduce the fungal burden (Figure 7A-D). The lack of prolonged survival for MMV689244 (EPL-BS1246) was surprising since it was the compound with the most potent *in vitro* activity after the reference compounds. Although overall survival was not prolonged, a reduction in grain number and grain mass was obtained with MMV689244 (EPL-BS1246). Since *M. mycetomatis* forms grains *in vivo* but not *in vitro,* we therefore hypothesized that compound MMV689244 (EPL-BS1246) might not be able to reach its target in the grain. We therefore tested the four fenarimol analogues with *in vitro* activity to determine if any would have activity against *M. mycetomatis* grains *in vivo.* It appeared that EPL-BS0178 (Log Rank p<0.0001), EPL-BS0495 (Log Rank, p=0.0199) and EPL-BS1025 (Log Rank, p=0.0436) were all able to prolong larval survival in a statistically significant manner (Figures 6E and 6F). The overall survival percentages of the three active fenarimol analogues were compared; treatment with analogue EPL-BS0178 resulted in a survival percentage of 36.7%, which was higher than that observed with analogues EPL-BS0495 (24.1%) and EPL-BS1025 (19.2%). Surprisingly, EPL-BS0178, EPL-BS0495 and EPL-BS1025 did not significantly lower the number of grains present in the infected larvae after three days of treatment (Table 2, Figure 7B). For EPL-BS1025, a significant reduction in total grain mass produced was noted (Figure 7D; median mass: 0.046 mm^2^; Mann-Whitney, p=0.042). The only fenarimol analogues which significantly lowered the total number of grains and the total grain mass were MMV689244 (EPL-BS1246) and EPL-BS0800 (median=18 grains; Mann-Whitney, p=0.019 and median=0.044mm^2^; Mann-Whitney, p=0.012) (Figures 7B and 7D), the two analogues which did not demonstrate enhanced larval survival. No difference in grain distribution (Table 2, Figure 8D, E, F) or larvae melanization was observed for any of the fenarimol analogues (Figure 7F).

To better understand the translation from *in vitro* activity to *in vivo* efficacy, physicochemical profiling of the ten compounds was performed. A correlation between the LogD at pH 7.4 (calculated using Stardrop [version 2017, Optibrium Ltd]), IC50 *in vitro* and percentage survival after 10 days was noted (Figure 9).

**Fig 9.**
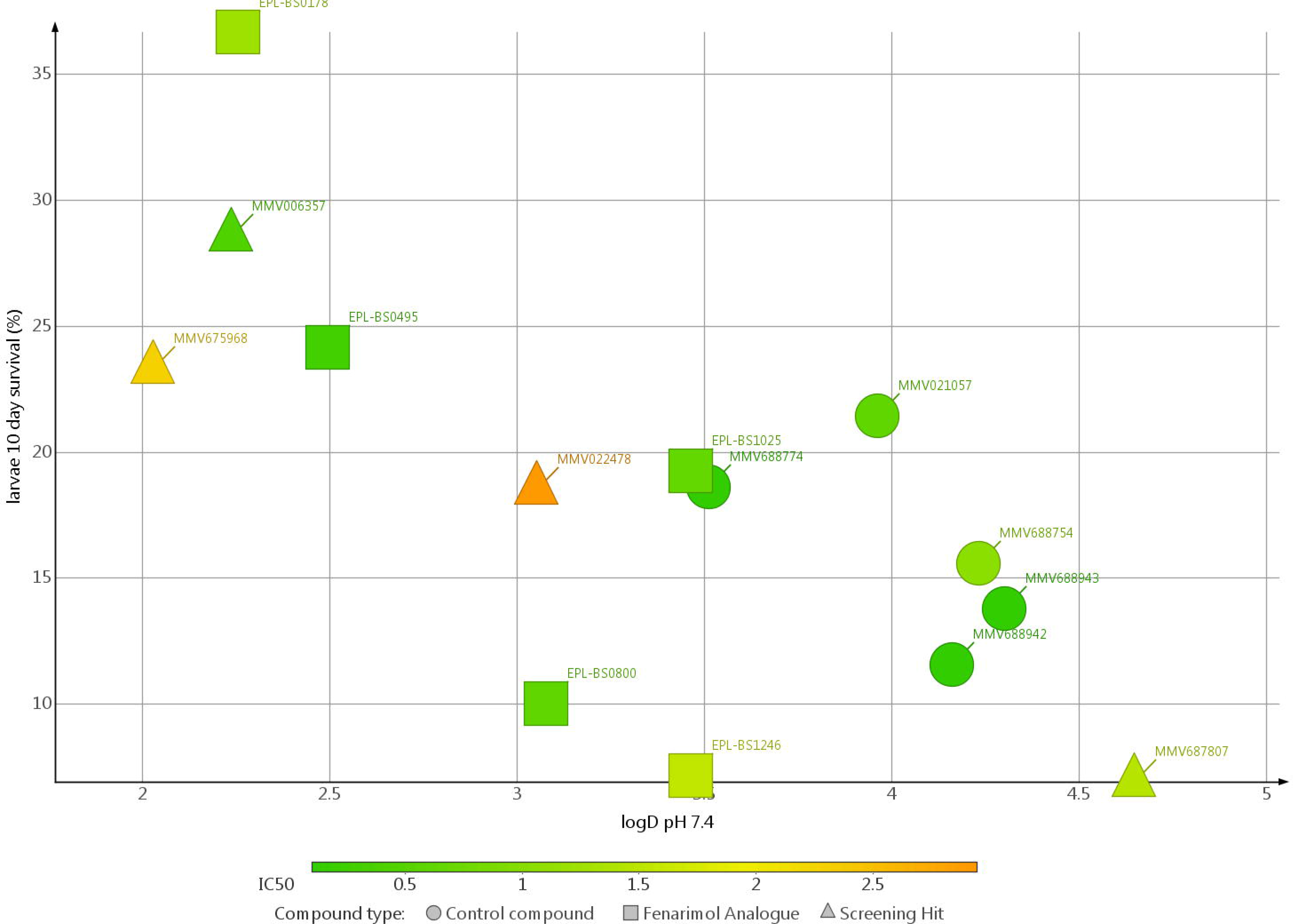
logD pH 7.4 versus *in vivo* larvae survival after 10 days

## Discussion

In this study, the Pathogen and Stasis Boxes were screened to identify compounds with *in vitro* antifungal activity against *Madurella mycetomatis.* Out of the 800 compounds tested, 13 compounds were associated with *in vitro* activity (determined as MIC50s) against *M. mycetomatis* strain Mm55 ranging from nanomolar (<0.007 µM) to micromolar (8 µM) potencies (S1 Table, Table 1, Figure 3). Of these 13 compounds, five are antifungal agents belonging either to the azoles (posaconazole, bitertanol and difenoconazole) or the strobilurins (azoxystrobin and trifloxystrobin), two are antiprotozoal agents (iodoquinol being an approved drug for the treatment of amoebiasis and MMV689244 a preclinical candidate for the treatment of Chagas disease) while the other five compounds had not previously been associated with any antifungal properties, being upstream exploratory molecules. Interestingly, four of the hits identified are known inhibitors of lanosterol 14αt-demethylase (or CYP51) - a cytochrome P450 enzyme involved in the conversion of lanosterol into ergosterol as part of the sterol biosynthesis pathway of fungi, yeast and trypanosomes - and a well-established antifungal drug target [32]. The antifungal activity of posaconazole against 34 clinical isolates of *M. mycetomatis* has previously been demonstrated [33]. In that study, an MIC50 of 0.06 µg/mL (0.086 µM) was obtained, which is 10 times higher than the MIC50 of <0.007 µM obtained in this study [33]. However, when we take only the seven strains tested in both studies in consideration, comparable MIC50s are obtained (MIC50 of <0.03 µg/ml (<0.04 µM) versus <0.007 µM) [33]. The antifungal activity of compounds MMV688774 (posaconazole), MMV688942 (bitertanol) and MMV688943 (difenoconazole) was also demonstrated when the Pathogen Box was screened with planktonic cells of other fungal species, such as *Candida albicans* [34] and *Cryptococcus neoformans* [34]. In that study, MMV688934 (tolfenpyrad) and MMV688271 were identified as antifungal agents [34], yet in our study MMV688934 did not inhibit *M. mycetomatis* growth at all, and MMV688271 only did so with an IC50 of 61.7 µM. In the same study, only six compounds were found to be able to inhibit *C. albicans* biofilm formation [35]. Among these compounds, amphotericin B, MMV687807, MMV687273 and MMV688768 were identified [35]. In our assay, amphotericin B had an IC50 of 9.7 µM and was therefore excluded from further evaluation, while MMV687807 was one of the 10 compounds with the highest *in vitro* activity.

*In vivo,* out of 10 compounds tested from the Pathogen and Stasis Boxes, five showed enhanced survival with the Log-Rank test (Figure 6). Of these compounds, only posaconazole had previously been tested in this larval model; larvae were treated with 5.7 mg/kg posaconazole, a dose similar to the one used in the treatment of human fungal infections, but this did not lead to any enhancement of larval survival [31]. In the current study, enhanced survival was noted when larvae were treated with 20 µM/larva which is comparable to 14 mg/kg of posaconazole. Furthermore, a reduction in the total number of fungal grains and total grain mass was noted. In this model, higher doses of posaconazole appear to be required to prolong larval survival, although there is a significantly greater than a thousand-fold disconnect between the concentrations required to demonstrate *in vitro* and *in vivo* activity. The compounds with the most potent *in vivo* activity were MMV006357, MMV675968 and MMV022478. MMV006357 was obtained from the Stasis Box and is a 2-aminothiazole derivative that was originally identified as a molecule with anti-mycobacterial activity [36, 37]. It has been demonstrated to be active against replicating and non-replicating mycobacteria and even had sterilizing activity against the latter [36], a property possessed by no other class of anti-mycobacterial compound. Furthermore, several of its analogues also had activity against drug-resistant isolates of *M. tuberculosis* [38]. In our *in vivo G. mellonella* model, this compound enhanced the overall survival significantly compared to that of the PBS treated group (Log-Rank, p=0.0012): the survival of 28.6% at day 10 recorded for MMV006357 was the highest observed with the compounds tested from the Pathogen and Stasis Boxes. However, the number of grains produced after three days of treatment and the total mass of these grains produced were not significantly reduced compared to the PBS treated group. This seems contradictory, but when counting grains in histology sections, we were not able to determine if these grains were still viable, and these results may indicate that a significant portion of the grains were killed but not cleared by the larval immune system. None of the compounds except MMV688943 (difenconazole) had a significant impact on melanization rate. As Eadie has described, compounds may influence the make-up of grains and their cement material, potentially increasing or decreasing the accessibility of *M. mycetomatis* to the larval immune system [39].

In addition to MMV006357, compounds MMV675968 and MMV022478 prolonged larval survival significantly. MMV675968 is a 2-4-diaminoquinazoline derivative active against *Cryptosporidium parvum* while MMV022478 is a pyrazolopyrimidine previously described as active *in vitro* against the asexual blood stage of *Plasmodium falciparum* as well as against *Plasmodium berghei* sporozoites (https://www.pathogenbox.org/). Although compounds MMV688943 (difenoconazole), MMV021057 (azoxystrobin), MMV688754 (trifloxystrobin), MMV689244 (EPL-BS1246) and MMV687807 had potent activity against *M. mycetomatis in vitro,* no enhanced survival was noted in infected *G. mellonella* larvae. Four of these five compounds, MMV688943 (difenoconazole), MMV021057 (azoxystrobin), MMV688754 (trifloxystrobin) and MMV689244 (EPL-BS1246) show a significant reduction in the total number of grains after three days of treatment. The mass of grains produced in mm^2^ was also significantly smaller than that of the PBS treated group. This indicates that some *in vivo* activity was displayed against the grain, but this did not in turn result in enhanced survival. For difenoconazole, this activity also resulted in a difference in grain size distribution in the larvae: fewer large grains and more small grains were produced compared to the PBS treated larvae. This might indicate that this compound is able to slow down grain formation and might be more effective when combined with other antifungal agents, or when administered prophylactically. The lack of association between *in vitro* susceptibility and *in vivo* efficacy against *M. mycetomatis* has been observed in the past, notably with respect to azoles. *In vitro, M. mycetomatis* was indeed shown to be most susceptible towards the azoles [26, 33] and less susceptible towards the polyene amphotericin B and the allylamine terbinafine [33]. However, only amphotericin B [26] and terbinafine, and not the azoles, were able to prolong *Galleria mellonella* larval survival [31]. This result was confirmed in a mouse efficacy model where amphotericin B was able to inhibit grain formation and itraconazole was not [40]. A lack of translation of activity from *in vitro* to *in vivo* could be due to pharmacodynamic considerations related to the mode of action of the drugs (i.e. speed of action, cidal versus static mechanisms) in the context of microorganisms being tested under significantly different *in vitro* and *in vivo* conditions (i.e. fast versus slow growth, high versus reduced metabolism). Another reason might be the difference in morphological organization of the fungus inside the host: while the fungal hyphae used for the *in vitro* testing are directly exposed to antifungal agents, *M. mycetomatis* forms grains *in vivo* in *G. mellonella* larvae – black, firm and brittle microstructures packed with fungal mycelia embedded in a hard brown-black cement-like material of still poorly understood nature [26, 27, 41, 42]. Azole antifungal agents are known to be more active on quickly dividing and faster growing fungi. *In vitro, M. mycetomatis* is metabolically active and visible growth is seen within days, while *in vivo,* grains are formed. Their metabolic activity is not currently known, but since no expansion of the grains is noted in tissue sections, it is envisioned that the grain itself is less metabolically active. Although lower metabolic activity within the grain might be one reason, it cannot be the only one. In earlier experiments performed by Murray, it was demonstrated that even 100 µg/mL of the fungicidal drug amphotericin B was not able to prevent *M. mycetomatis* growth when freshly isolated *M. mycetomatis* grains from infected mice were immersed in melted agar, while 8 µg/mL was enough to inhibit the growth of *M. mycetomatis* cultured mycelia [43, 44]. Although the grain itself is a key feature of mycetoma, little is known about its constituents or metabolic activity. In the 1970s some attempts were made by Findlay and Vismer to identify the constituents, but since then little progress has been made [45–47]. Findlay showed that the grain owed its existence and its toughness to a tanning of the structural and inflammatory proteins of the host by a diffusible melanin-type pigment synthesized and secreted by the fungus [45]. When this melanin was isolated and added to our *in vitro* susceptibility assay, MICs for ketoconazole and itraconazole were elevated 32 times [48]. Within the grain, intrahyphal growth is observed [47] and the hyphae themselves are embedded in cement material, which makes it difficult for each drug to reach the metabolically active part of the fungus. Although the exact constituents of this cement material are still unknown, chitin [49] and beta-glucan [50] are known to be involved. These constituents are implicated in the reduced antifungal susceptibility of fungal biofilms [51]. In addition, the grain itself is surrounded by a collagen capsule, which also makes it more difficult for the drug to reach the grain itself [52].

We postulated that the activity against the fungal grain of the hits identified *via in vitro* and *in vivo* screening might be enhanced by chemical modification of the original hit structure. This hypothesis was first validated with respect to hit MMV689244 (EPL-BS1246), one of the most potent *in vitro* hits identified during screening of the Pathogen Box but which did not, however, significantly enhance survival when evaluated in the *G. mellonella* larvae model. MMV689244 is a fenarimol analogue, identified as a potent inhibitor of *Trypsonosoma cruzi* during a targeted screening exercise for new drugs for Chagas disease [15–17], during which over 800 fenarimol analogues were synthesized. By screening 35 fenarimol analogues representative of the chemical diversity of this scaffold, we identified an additional four fenarimol analogues with *in vitro* activity against *M. mycetomatis:* EPL-BS0178, EPL-BS0495, EPL-BS0800 and EPLB-BS1025. Three of these fenarimol analogues (EPL-BS0178, EPL-BS0495 and EPL-BS1025) showed potent *in vivo* activity (Figure 6), further validating the chemotype as an antifungal agent. Of these fenarimol analogues, EPL-BS0178 gave the longest overall survival percentage, with an overall value of 36.7% at day 10 (Log-Rank, p<0.0001), the highest overall survival ever obtained in this model. However, once again, this compound did not reduce the total number of grains present or their total mass. Additionally, none of the fenarimol analogues had any impact on melanization rate. Interestingly, there appears to be a correlation between the polarity (expressed as logD at pH 7.4, see Figure 9) of the hit compounds and the prolongation of survival in the *G. mellonella* larval assay: across the few chemotypes investigated, those compounds with logD values>2.5 were the best performers in this *in vivo* assay model. It is therefore proposed that potency against *M. mycetomatis in vivo* is determined by a play off between the general antifungal activity and the physicochemical properties of the compound, and that this needs to be taken into consideration when seeking future analogues and antifungal agents against *M. mycetomatis* [53]. In light of these findings, it would be interesting to determine if other fenarimol analogues or analogues of 2-aminothiazole increase the therapeutic efficacy against *M. mycetomatis* grains *in vitro* and eventually *in vivo.* At present, over 765 of the original fenarimol analogues and a number of 2-aminothiazole analogues series synthesized by other groups have yet to be screened [37, 38]. With these important new results in hand, we propose opening the next stages of this project up to the wider community by adopting an Open Source approach that has previously resulted in effective research consortia in antimalarial drug discovery [54]. We have deposited the data associated with this work in an online database (http://tinyurl.com/MycetomaMols). have started an online discussion area on two websites to gather community expertise (https://github.com/OpenSourceMvcetoma and https://www.reddit.com/r/OpenSourceMvcetoma/), have opened up the electronic laboratory notebook associated with the chemical resynthesis of fenarimol analogs and have started a social media account for community use and outreach (https://twitter.com/MycetOS). We have used these sites to clarify the current needs of the community, which are i) samples of analogs of the most promising compounds described in this paper and ii) advice on which of the existing compounds to which we have access should next be evaluated *in vitro* and *in vivo;* some directions in the fenarimol library have been suggested (Figure 10). Gathered together, these resources constitute *Open Source Mycetoma* (MycetOS), a project that will adhere to six basic laws of open source research, most importantly that all data and ideas are freely shared, that anyone may participate and that there will be no patents [54]. The authors of this paper, and DND*i* itself, are partners in such an approach, but any other member of the community is free to participate and contribute as an equal partner provided the principle of open work is upheld. We hope that this initiative, coupled with the promising new hits we have reported, will lead to progress in drug discovery for this most neglected of neglected tropical diseases.

**Fig 10.**
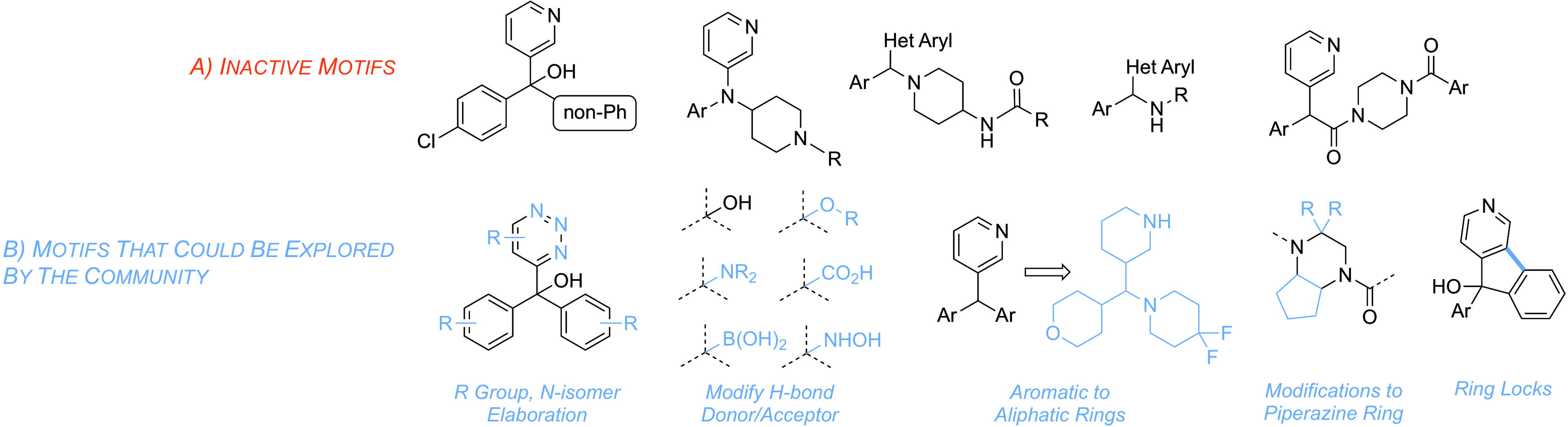
Summary of inactive motifs discovered by screening compounds from the Epichem library (A) and potential motifs for future exploration by the open source community (B).

## Acknowledgement

The authors are indebted to Medicines for Malaria Venture (MMV) for providing both the Pathogen Box and the Stasis Box free of charge. The fenarimol collection was made available by DNDi. The synthesis of this collection of compounds was performed by Epichem and funded by DNDi.

## Supporting Information Legends

S1 Table. Information and data of all compounds tested from the Pathogen Box, Stasis Box and fenarimol analogues.

S2 Text. General method for chemical synthesis of hit compounds and novel analogues.

